# SARS-CoV-2 cell-to-cell infection is resistant to neutralizing antibodies

**DOI:** 10.1101/2021.05.04.442701

**Authors:** Natalia Kruglova, Andrei Siniavin, Vladimir Gushchin, Dmitriy Mazurov

**Author notes:** Correspondence should be addressed to D.M.

## Abstract

The COVID-19 pandemic caused by SARS-CoV-2 has posed a global threat to human lives and economics. One of the best ways to determine protection against the infection is to quantify the neutralizing activity of serum antibodies. Multiple assays have been developed to validate SARS-CoV-2 neutralization; most of them utilized lentiviral or vesicular stomatitis virus-based particles pseudotyped with the spike (S) protein, making them safe and acceptable to work with in many labs. However, these systems are only capable of measuring infection with purified particles. This study has developed a pseudoviral assay with replication-dependent reporter vectors that can accurately quantify the level of infection directly from the virus producing cell to the permissive target cell. Comparative analysis of cell-free and cell-to-cell infection revealed that the neutralizing activity of convalescent sera was more than tenfold lower in cell cocultures than in the cell-free mode of infection. As the pseudoviral system could not properly model the mechanisms of SARS-CoV-2 transmission, similar experiments were performed with replication-competent coronavirus, which detected nearly complete SARS-CoV-2 cell-to-cell infection resistance to neutralization by convalescent sera. Based on available studies, this is the first attempt to quantitatively measure SARS-CoV-2 cell-to-cell infection, for which the mechanisms are largely unknown. The findings suggest that this route of SARS-CoV-2 transmission could be of great importance for treatment and prevention of COVID-19.

**Importance:** Immune surveillance of viral or bacterial infections is largely mediated by neutralizing antibodies. Antibodies against the SARS-CoV-2 spike protein are produced after vaccination or infection, but their titers only partly reflect the degree of protection against infection. To identify protective antibodies, a neutralization test with replicating viruses or pseudoviruses (PVs) is required. This study developed lentiviral-based PV neutralization assays that, unlike similar systems reported earlier, enable quantitative measurement of SARS-CoV-2 neutralization in cell cocultures. Using both PVs and replication-competent virus, it was demonstrated that SARS-CoV-2 cell-to-cell infection is considerably more resistant to serum neutralization than infection with purified viral particles. The tests are easy to set up in many labs, and are believed to be more informative for monitoring SARS-CoV-2 collective immunity or entry inhibitor screening.

## Introduction

SARS-CoV-2 is a respiratory virus, a causative agent of COVID-19. The primary target of the virus is the airway epithelium of the upper respiratory tract.^1–3^ During the course of the disease, the virus can descend to the lower respiratory tract, infecting bronchial epithelium and type II pneumocytes.^4^ The main receptor for SARS-CoV-2, angiotensin-converting enzyme 2 (ACE2),^1,5,6^ determines the viral tropism, which is not restricted to the respiratory epithelium and in certain cases can infect enterocytes, as well as kidney, heart, brain, and other cell types.^7,8^ Molecules other than ACE2 have been reported to be involved in SARS-CoV-2 entry, such as neuropilin-1,^9,10^ AXL,^11^ and CD147, although the role of the latter is speculative.^12,13^

SARS-CoV-2 entry is mediated by the spike (S) protein.^14^ The S protein belongs to trimeric class I fusion proteins^15^ that undergo substantial conformational changes when bound to a cellular receptor, leading to fusion between viral and cell membranes.^16,17^ The extracellular portion of the spike consists of two subunits: S1 binds to ACE2 and S2 mediates the viral fusion.^17^ A newly synthesized spike exists in a metastable prefusion state.^17^ Following attachment to permissive cells, the receptor binding domain (RBD) in the S1 subunit transitions between the inactive ‘down’ position and the accessible ‘up’ position for interaction with ACE2.^18–20^ However, binding the S protein to ACE2 is not sufficient for triggering membrane fusion, because the fusion peptides of coronaviral S proteins have a ‘hidden’ localization inside the S2 subunit.^17^ Proteolytic cleavage at the S2’ site releases fusion peptide. This process is mediated by several host proteases: TMPRSS2, and lysosomal cathepsins B and L.^6,21^ Depending on localization within the target cells, these proteases largely determine virus entry sites; plasma membrane in the case of TMPRRS2 or endosomes when cathepsins are engaged.^14,22^ In this regard, SARS-CoV-2 is not unique and demonstrates features that have long been known from other coronaviruses.^21^ In contrast to the S2’ site, the furin cleavage site at the S1/S2 boundary is a special feature of SARS-CoV-2 that generally distinguishes it from other beta-coronaviruses, such as SARS-CoV,^23^ with the exception of MERS-CoV, where it is present.^21^

An invaluable instrument for coronavirus entry inhibitor assessment is pseudoviruses. They are safe, reliable, and fast for generating quantitative data relative to fully competent viruses, which often require strict regulation when working with them. The SARS-CoV-2 spike protein is sufficient to mediate pseudovirus entry and many pseudoviral systems were developed during the COVID-19 pandemic, generally using HIV,^6,24–28^ MLV,^2,29,30^ or VSV^3,5,31,32^ platforms. In comparison to retro- or lentiviral particles, which require 48 hours to get infectivity results, results for VSV-based particles can be obtained within 24 hours of infection and at higher titers, although the production is more labor-intensive.^6,33,34^ In general, the choice of pseudoviral system is primarily dictated by the preferences of a particular research group.^35^ Using pseudoviral tests, large amounts of data on the inhibitory activity of sera from convalescent and vaccinated individuals, monoclonal antibodies, proteins, peptides, and small molecules were collected and analyzed.^25–29,36^

Despite fast progress in SARS-CoV-2 entry inhibitor evaluation using pseudoviruses, the vast majority of developed systems are capable of measuring infectivity only with purified particles. Meanwhile, the largely unknown – and potentially important – mechanism of SARS-CoV-2 spread from cell-to-cell has not been evaluated with pseudoviruses. This study describes the S protein pseudotyped lentiviral system for measuring SARS-CoV-2 infection in both cell-free and cell-to-cell infection settings. This was achieved with replication-dependent reporter vectors that were developed earlier.^37,38^ The key feature of these vectors is that the reporter is silent in the pseudovirus-producing cells, but active after infection of the target cells and completion of one cycle of viral replication. This enables infectious events to be measured directly in cocultures of producer and target cells at zero background level. The concept was effectuated by placing a reporter cassette in reverse orientation relative to HIV-1 genomic RNA and through interruption it with an intron, that prevented a functional reporter protein expression from LTR and CMV promoters in transfected (producer) cells. Comparative analysis of SARS-CoV-2 infection in two transmission settings revealed a substantially lower capacity of convalescent sera to neutralize infection in cell cocultures than in a cell-free test. This effect was reproduced with replication-competent SARS-CoV-2, indicating that cell-to-cell transmission of SARS-CoV-2 and its elevated resistance to entry inhibitors are important parameters for monitoring anti-viral immunity and developing anti-coronaviral drugs.

## Results

### Generation and optimization of a SARS-CoV-2 pseudoviral system to measure cell-free infection

Pseudoviruses (PVs) are viruses enveloped with a heterologous surface protein that changes their natural tropism. Unlike native systems, heterologous protein envelopes are often incorporated into PV particles at lower efficiency. A number of studies have been focused on optimizing lenti- and retroviral systems pseudotyped with the SARS-CoV S protein. Giroglou et al. showed that C-terminal truncation of the S protein increased retroviral particle titers, explained by the removal of the ER retention signal and exposure of the S protein to the cell surface.^39^ Moore et al. found that both C-terminal truncation and substitution of the cytoplasmic portion of the S protein with eight amino acids from the C-terminus of HIV-1 gp41 increased the level of infectious lentiviral particle production.^40^ Later on, the spike protein from SARS-CoV-2 with a cytoplasmic portion deleted was used in a number of pseudoviral test systems.^6,32,41–43^ The furin cleavage site that was present in SARS-CoV-2 S but not in the SARS-CoV S protein is thought to be involved in spike maturation, virus entry, and syncytium formation^5^ and, therefore, can also affect infectivity measured with PVs.

In order to establish an HIV-based infection system, the SARS-CoV-2 S protein was modified by deleting the last 19 amino acids (ΔC19) or substituting them with eight amino acids from HIV-1 gp41 (H2). These modifications were either combined with the mutation in the furin cleavage site RRAR to A (ΔF), or left uncombined, to generate the six variants of spike protein indicated in Figure 1A. Next, a SARS-CoV-2 permissive HEK 293T cell line was established with a stable expression of the human ACE2 receptor via lentiviral transduction and FACS sorting (Figure 1B). A cell-free infectivity assay was set up, as schematically illustrated in Figure 1C. PVs were generated by co-transfecting 293T cells with one of the S protein coding plasmids, HIV-1 packaging vector pCMV-dR8-2, and an improved intron-regulated reporter vector pUCHR-inLuc-mR, capable of measuring both cell-free and cell coculture infections using the mean of luciferase activity.^37,38^ Additionally, the pUCHR-IR-GFP reporter plasmid without an intron was used to evaluate cell-free infectivity levels using flow cytometry. At 48 hours post-transfection, supernatants containing PVs were harvested and concentrated by centrifugation. Equal amounts of PVs were added to 293T/ACE2 cells for 48 hours, and levels of infection were estimated by measuring luciferase activity or percentage of GFP-positive cells, depending on reporter type. The resulting values of infection were normalized to p24 levels, and presented relative to the values obtained for the wild-type S protein. As shown in Figure 1D, ΔC19 moderately increased the level of infection, while the H2 modification had no or little effect on infectivity. By contrast, the ΔF mutation resulted in about a 1.5 log increase in PV infectivity. On ΔF background, however, the improving effect of ΔC19 was much less pronounced than detected without ΔF. PV titration was used to confirm a substantial effect of the ΔF mutation on the level of PV transduction (∼20 fold enhancement in many PV dilutions) (Figures 1E and F). The increased infectivity of the ΔF mutant PVs was not accompanied by an increase in S protein expression on PV-producing 293T cells (Figure 1G). Thus, it was unclear whether ΔF infectivity was enhanced from S incorporation into PVs or if this was a feature of the 293T cellular system, in which S processing by furin is important during the fusion step of the viral life cycle.

**Figure 1.**
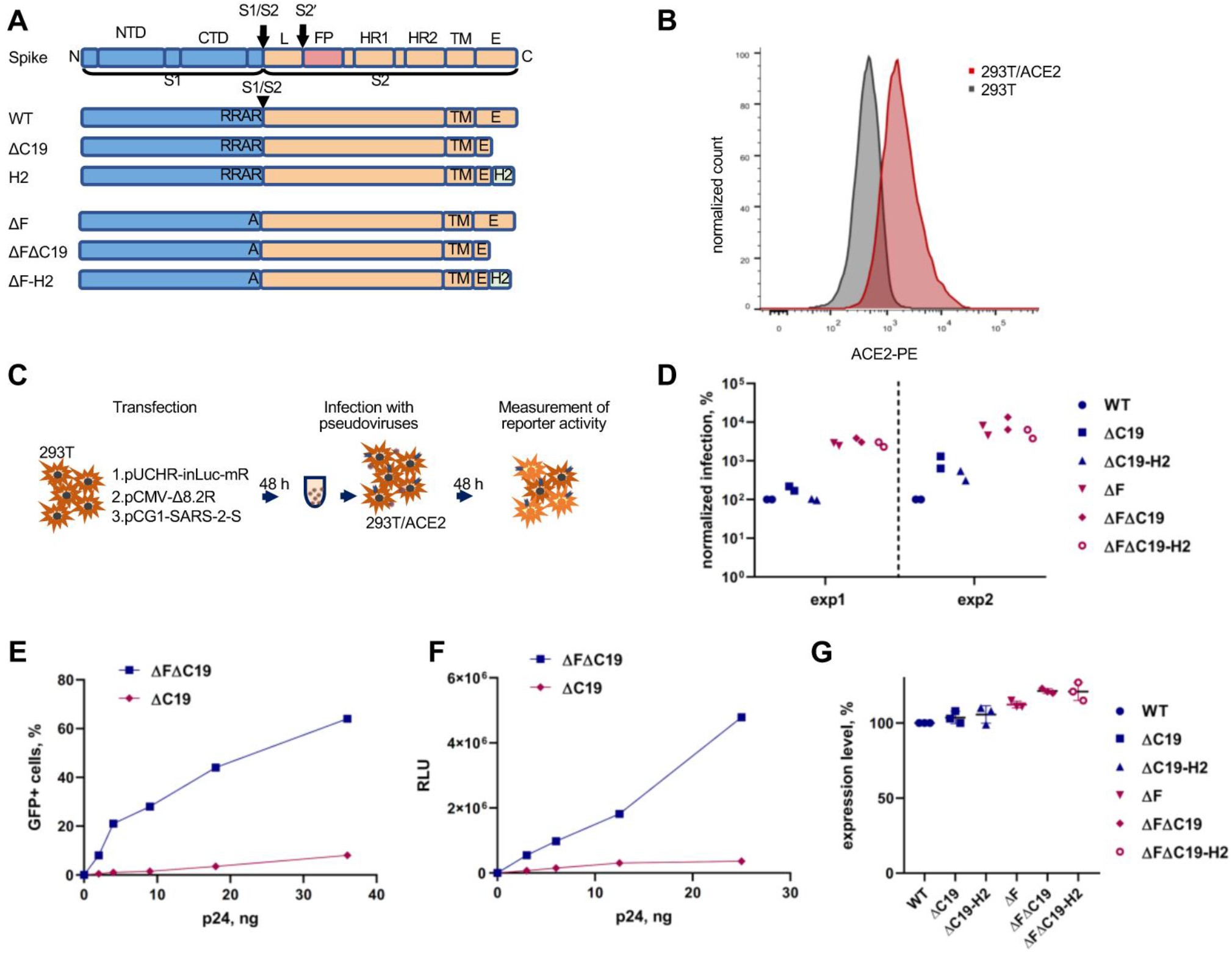
Development of a SARS-CoV-2 cell-free infection test with PVs. **A**. A schematic illustration for S-protein variants used in pseudovirus infection tests. Six different constructs of the S protein were generated by PCR mutagenesis **B**. Evaluation of the ACE2 surface expression on 293T cells stably transduced with the hACE2 using flow cytometry. **C**. Experimental setup for SARS-CoV-2 cell-free infection measurement. **D**. The levels of Infection detected with different variants of the S protein. PVs were added to 293T/ACE2 cells in a equal amount based on HIV-1 Gag quantification. The luciferase activity measured for a mutant spike was normalized to that obtained for the wild-type S protein. Two independent experiments with two different PV preparations were performed. **E**,**F**. The levels of cell-free infection with indicated PVs were measured using either GFP (**E**) or inLuc (**F**) reporter. **G**. The levels of S protein expression on PV producing cells estimated by flow cytometry. 293T cells were transfected to express indicated variants of protein S and stained with convalescent human serum in 48 hours. Median fluorescence intensity (MFI) level was calculated for the every mutant in the gate of transfected cells and normalized to the MFI detected for wild-type S protein.

In summary, a SARS-CoV-2 cell-free infection test was developed in a 24-well plate format with a high level of sensitivity. Using the ΔFΔC19 modification of the SARS-CoV-2 S protein, 50-60% GFP transduction and about 4 log over the background elevation of luciferase activity was achieved, making consecutive inhibitory analysis accurate and reproducible.

### Development of a SARS-CoV-2 pseudoviral system to quantify cell-to-cell infection

In order to evaluate SARS-CoV-2 cell-to-cell infection, a one step transfection-infection assay with the inLuc-mR reporter vector described earlier was set up.^37,38^ Briefly, 293T/ACE2 cells were co-transfected with viral vectors, as outlined above, for cell-free infection. In approximately 12-16 hours, transfected cells started to produce PVs, which infected nearby 293T/ACE2 cells. At 48 hours post-transfection, one cycle of replication was complete and luciferase activity can be measured (Figure 2A). Using this assay, the levels of infection with one of the six variants of the S protein were quantified. Wild type, ΔC19, and H2 proteins did not mediate infection at all, however, all three variants bearing the ΔF mutation supported a good level of infection. The addition of ΔC19 to ΔF increased the level of infectivity by 0.5 log, while the H2 modification had no effect on the signal (Figure 2B). We have previously demonstrated that intron-regulated reporter vectors do not detect infection in cell syncytia, as the reporter protein can be expressed only in actively replicating target cells.^37^ Therefore, the ability of differently modified S protein variants to induce syncytia formation in 293T/ACE2 cells was examined. Consistent with previously reported data,^6,22,36^ a massive cell-cell fusion upon expression of all three variants of spike bearing the furin cleavage site was detected, and there was no syncytia formation in the samples transfected with ΔF variants (Figure 2C). This suggests that S-mediated syncytia formation inhibits lentiviral reporter expression in permissive cells; consequently, wild type S protein cannot be used to assess cell-to-cell infection in the 293T/ACE2 cellular model. In summary, the possibility of measuring SARS-CoV-2 cell-to-cell infection using the intron-regulated luciferase vector was demonstrated, and the ΔFΔC19 mutant of S was selected for further study.

**Figure 2.**
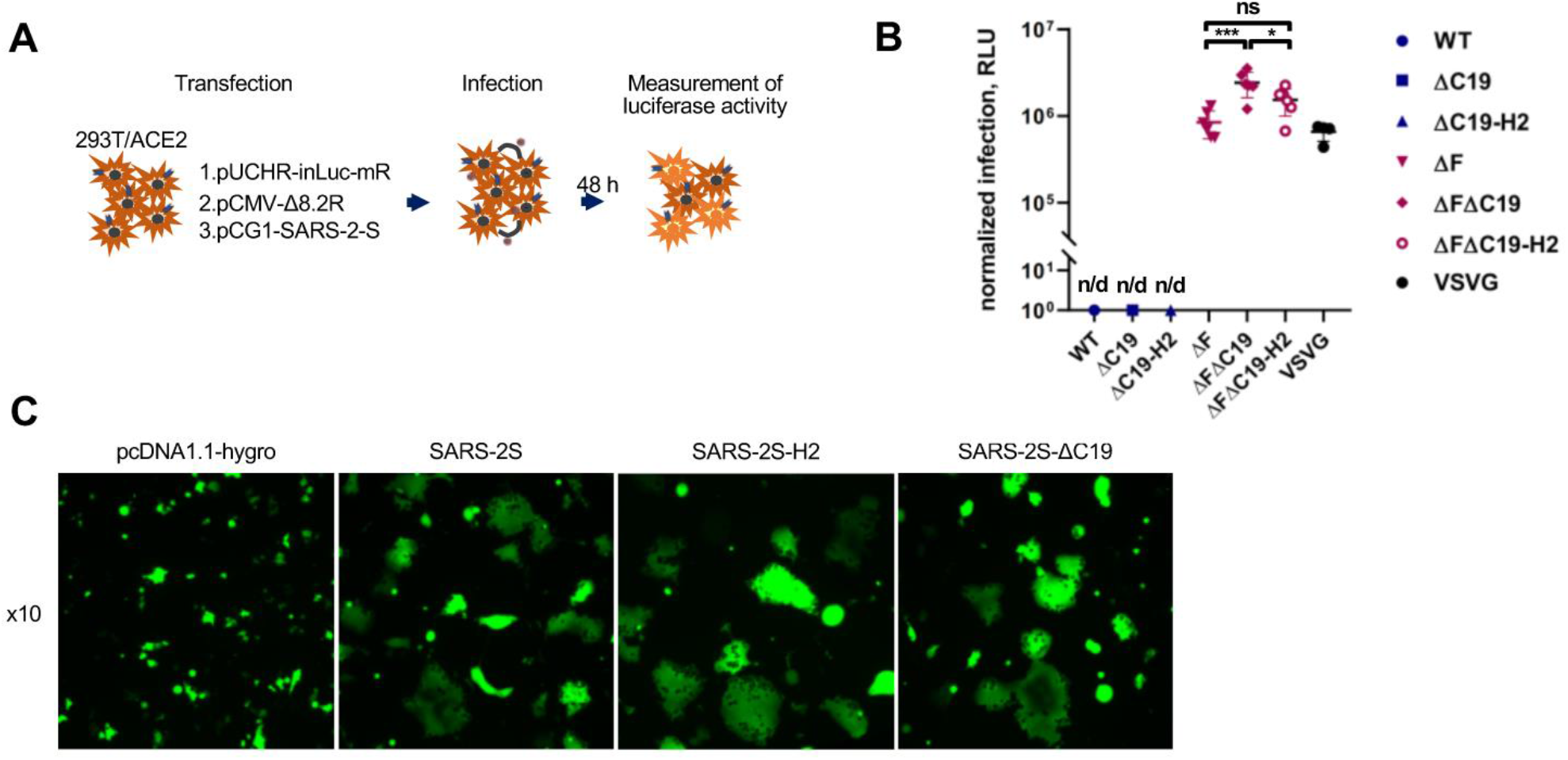
Establishing PV system for measurement of SARS-CoV-2 cell co-culture infection. **A**. A schematic representation of SARS-CoV-2 one step transfection-infection assay in 293T/ACE2. **B**. The levels of infectivity measured using one step assay with different spike protein mutants. Samples with the intact furin cleavage site produced no signal above background (n/d). The differences between ΔF mutants were calculated by one-way ANOVA with the Tukey’s multiple comparison test, and are significant at p=0.0008(***) and p=0.0457 (*). **C**. Syncytia formation induced by wild-type or mutant SARS-CoV-2 S protein. 293T cells were co-transfected with plasmids for expression of GFP and one of indicated variant of S protein with the intact furin cleavage site. At 24 hours posttransfection, cells were detached with 1 mM EDTA and mixed with 293T/ACE2 cells at 1:1 ratio for another 24 hours. Typical images of cells captured on epifluorescent microscope with a filters set for FITC are demonstrated.

### Comparative analysis of the neutralizing activity of convalescent sera in SARS-CoV-2 cell-free and cell coculture pseudoviral infection tests

Using the developed pseudoviral infection tests, side-by-side comparisons of the neutralizing activity of convalescent sera from COVID-19 patients in cell-free and cell coculture modes of infection were performed. To avoid possible biases that can be observed when a neutralizing agent is added at the time of infection initiation, neutralization tests were designed to allow either PVs or producer cells to be preincubated with a serum for 1 hour prior to the target cell addition (see schematic in Figure 3A). Specifically, cell-free PVs in the amount of 10 ng of p24 were incubated with indicated serum dilutions in a total volume of 400 µl of culture medium, and added to 8×10^4^ 293T/ACE2 cells, seeded overnight in a 24-well plate. The levels of cell-free infection were estimated 48 hours later by measuring luciferase activity in cell lysates. In these experimental settings, the results with control samples were consistently reproduced at the level of ∼10^6^ RLU, giving an opportunity to detect a wide range of inhibitory activity. Five COVID-19 convalescent sera with high neutralizing activity were selected and evaluated in the cell-free infection test with ΔFΔC19. As shown in Figure 3A and E, all samples demonstrated NT_50_ in a range between 1/1500 and 1/12000 dilution, whereas a non-immune serum had no inhibitory activity. Additionally, in order to determine whether the furin cleavage site mutation influenced neutralization titer, ΔC19 variant *per se* was tested. The inhibition rates against ΔC19 were slightly higher than those for ΔFΔC19 (Figure 3B), including NT_50_ values (Figure 3C). Nevertheless, similar titration curves for all tested sera were observed with both ΔFΔC19 and ΔC19. Thus, the ΔF mutation did not dramatically change S protein neutralization in the cell-free test, but was absolutely necessary for measuring cell coculture infectivity and making the correct comparison between two types of infection.

**Figure 3.**
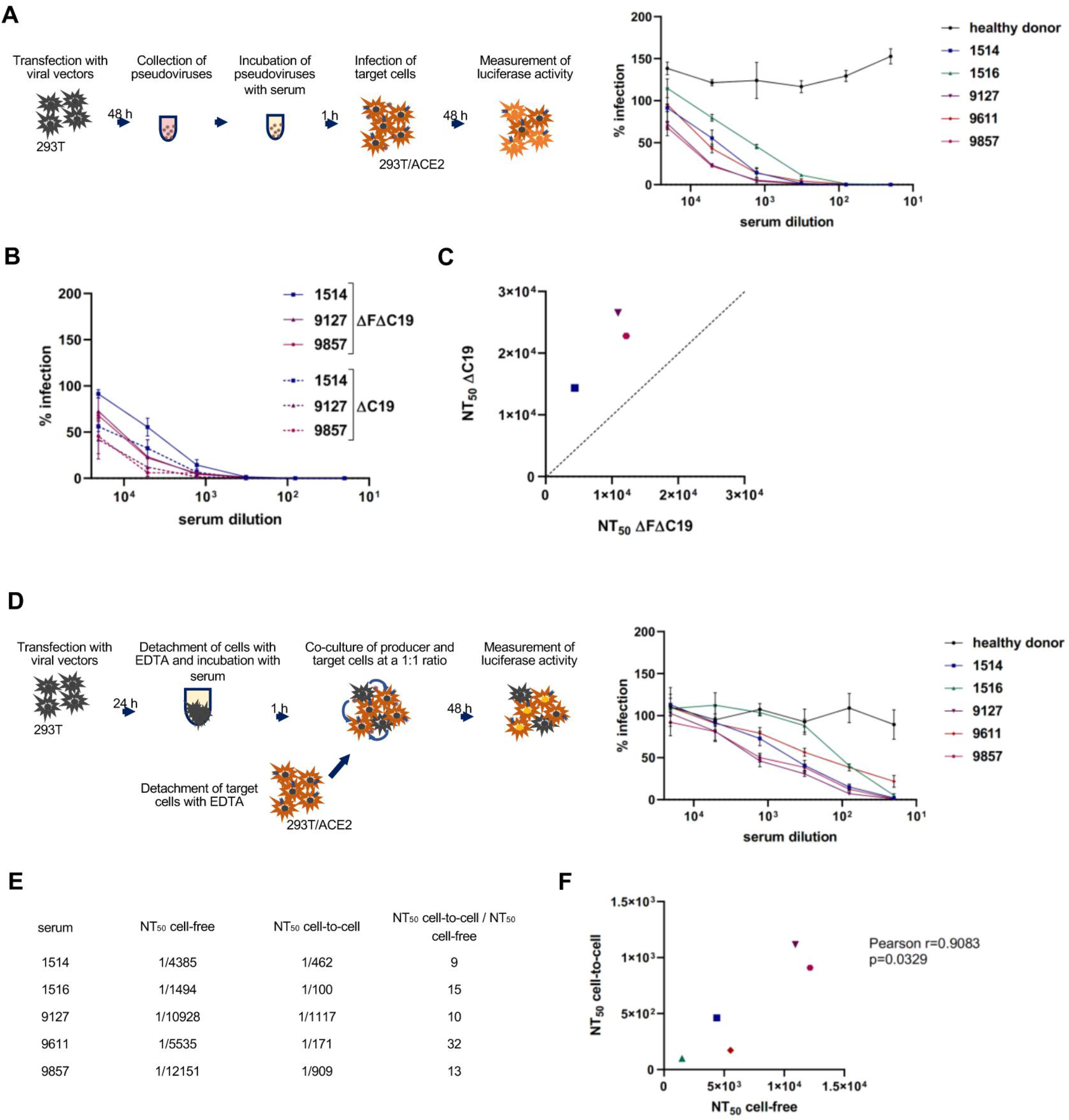
Neutralization activity of convalescent sera determined using SARS-CoV-2 PVs. **A**. The experimental steps designed for cell-free neutrilization test. Viral particles pseudotyped with ΔFΔC19 were preincubated with human serum dillution for 1 h and added to the 293T/ACE2 target cells. The control RLU values obtained without serum were set at 100%. The levels of infection detected in the presence of immune or non-immune serum were presented relative to control.. **B**. Neutralizing activity of convalescent sera against two indicated S protein mutants measured in a cell-free infection test. The assay was set up as in A. **C**. Correlation between 50% serum neutralizing titers (NT_50_) obtained against SΔFΔC19-PVs and SΔC19-PVs. **D**. A schematic illustrating cell coculture neutrilization assay setup (on the left) and neutrilization curves (on the right) obtained for indicated sera in this test. 293T cells transfected with viral vectors for 24 hours were detached with 1 mM EDTA, incubated with a serially diluted serum for 1 h, and co-cultured with 293T/ACE2 cells at 1:1 ratio for 48h. Data were collected and presented as in A. The average results from three independent experiments ± standard deviations are shown in A, B and D. **E**. Comparison of the 50% serum neutralizing titers (NT_50_) obtained in cell-free and cell coculture infection tests with PVs. The values were extracted from data presented in A and D. **F**. The correlation between cell-free and cell coculture neutralizing titers detected for five convalescent sera.

The cell-to-cell neutralization test was designed to be as similar as possible to settings used for the cell-free PV inhibition analysis. To generate SARS-CoV-2 producer cells, non-permissive 293T cells were co-transfected with pCMV-Δ8.2R, pUCHR-inLuc-mR, and pCG1-SARS-2-SΔFΔC19 plasmids, as described for cell-free infection. After 24 hours, cells were gently suspended using ethylenediaminetetraacetic acid (EDTA) and washed once with phosphate buffered saline (PBS); 2.6×10^4^ transfected cells in 200 µl culture medium were preincubated with a certain serum dilution for 1 hour at +4°C and mixed with 5.6×10^4^ 293T/ACE2 cells, giving a total of 8×10^4^ cells in 0.4 ml of culture medium. The cell mixture was placed in the wells of a 24-well plate and incubated for 48 hours before luciferase activity measurement (Figure 3D, schematic). The described format, and the resulting ratio of one producer cell to two target cells, provided the optimal sensitivity for measuring cell coculture infection and comparing it to cell-free infection in control samples. As shown in Figure 3D on the right, inhibition of SARS-CoV-2 cell coculture infection required high concentrations of convalescent sera, with NT_50_ detected within the 1/100 to 1/1100 dilution range. Compared to the serum activities against cell-free infection, the neutralization capacities of the same sera against cell coculture infection were more than tenfold lower (Figure 3E). Nonetheless, NT_50_ titers of individual serum samples measured in cell-free and cell-to-cell infection tests correlated to each other (Figure 3F), i.e., sera with a higher inhibitory titer detected in the cell-free infection test more effectively inhibited cell coculture infection.

In summary, by using the developed pseudoviral single cycle replication assay with the intron-regulated reporter vector, it was demonstrated that SARS-CoV-2 cell coculture infection was much more resistant to neutralization by convalescent sera than infection with purified PVs.

### Neutralization potential of convalescent sera against replication-competent SARS-CoV-2

Neutralization tests with PVs, although safe, have serious limitations, since they can only mimic the entry step of the viral life cycle. The mechanisms of viral assembly, egress, and transmission for the HIV and coronaviruses are very different, so the HIV-1 core proteins responsible for these processes – and used in this study’s pseudoviral tests – cannot model SARS-CoV-2 cell-to-cell transmission. With an understanding of all the drawbacks of the developed tests, an investigation of whether the resistance of cell-to-cell transmission to antibody neutralization could be reproduced with a full replication-competent SARS-CoV-2 was conducted. To this end, Vero E6 monkey fibroblast cells were chosen for virus production, setting up cell-free and cell-to-cell infection. The 293T/ACE2 cells for this purpose were excluded, as they died quickly after infection with coronavirus, making viral stock generation or maintaining multiple cycles of replication impossible. To remain consistent with pseudoviral tests, the number of plated Vero cells were proportionally similar to what was used for 293T cells. Neutralization of cell-free infection was performed by preincubating 0.01 MOI of SARS-CoV-2 strain hCoV-19/Russia/Moscow_PMVL-4 with serially diluted convalescent sera for 1 hour, and then adding to the Vero cells, seeded in a 96-well plate overnight. At day five post infection, cytopathic effect (CPE) was measured using the MTT test. As shown in Figure 4A, all sera completely blocked SARS-CoV-2 replication at 1/100 dilution; NT_50_ values ranged from 1/400 to 1/1400. These values were lower than the corresponding NT_50_ determined in the pseudoviral test. This can be explained by the doses of PVs and virions used for the neutralization assays, which are difficult to compare or normalize. Nevertheless, the results of two cell-free assays correlated well to each other (Figure 4B).

**Figure 4.**
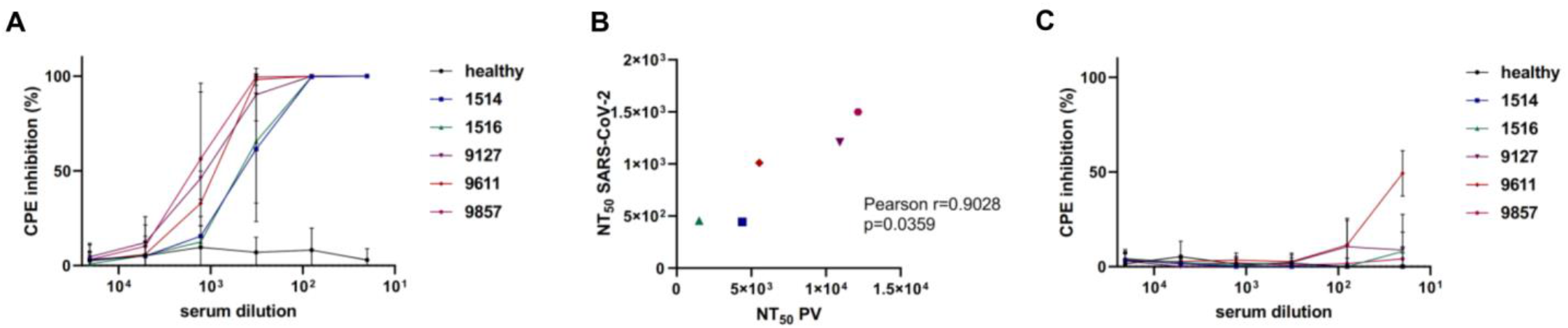
Neutralization activity of convalescent sera against replication-competent SARS-CoV-2. **A**. Serum activity against the purified SARS-CoV-2 virions. Serial fourfold dilutions of serum samples from convalescent donors were incubated with SARS-CoV-2 and added to the VeroE6 cells. Cell survival was determined 5 days later by MTT test and expressed as a percentage of cytopathic effect (CPE) inhibition measured relative to untreated control, which was set at 100%. **B**. Correlation between NT_50_ serum neutralizing titers measured with the ΔFΔC19-PVs and the live SARS-CoV-2. **C**. Serum neutrilizing activity against SARS-CoV-2 in cell coculture conditions. Serial fourfold dilutions of serum samples from convalescent donors were incubated with SARS-CoV-2 infected VeroE6 cells, which were then mixed with uninfected VeroE6 cells at 1:1 ratio. The results were collected and presented as in A. The average results from three independent experiments with standard deviations are shown in A and C.

Cell-to-cell SARS-CoV-2 spreading assay was initiated by infecting 2.6×10^4^ Vero E6 cells with fully competent virus for 24 hours, followed by PBS washing and preincubating with a serum before it was added to 5.6×10^4^ uninfected Vero E6 cells. The level of CPE was measured five days later using the MTT test. In stark contrast to the cell-free infectious test, the majority of the serum samples, even at minimal dilution, did not prevent cytopathic effect of fully-competent SARS-CoV-2 (Figure 4C), with the exception of serum 9611, which at 1/20 dilution displayed ∼50% inhibitory activity.

In conclusion, it was demonstrated that cell-to-cell spread of the fully competent wild type SARS-CoV-2 is almost completely resistant to convalescent serum neutralization. This effect was even more pronounced than the resistance detected using pseudoviruses.

## Discussion

The fast and global spread of COVID-19 requires unprecedent efforts to control this pandemic. An important parameter of collective immunity, gained after SARS-CoV-2 infection or vaccination, is the neutralization activity of anti-spike antibodies. This reflects the degree to which the studied population is protected from the infection, and provides more adequate information than a titer of anti-spike antibodies measured by ELISA. However, this test requires strict BSL3 conditions in order to work with highly pathogenic full-length SARS-CoV-2, and it has not been widely utilized. In this respect, different PV systems are considered safe and acceptable for many labs, since they allow the completion of only one cycle of viral replication. The PVs have been adapted by many researchers to characterize the SARS-CoV-2 entry process,^3,5,6,22,44,45^ monitor the dynamics of neutralizing humoral immunity to SARS-CoV-2,^14,46,47^ and screen potential inhibitors.^27,36^

The aim of this study was to not just replicate a PV system but rather, develop a lentivirus-based PV test capable of measuring SARS-CoV-2 infection, both with purified PVs and in cell cocultures. The latter has not been appreciated previously or measured accurately. The use of identical vectors to initiate both types of infection makes comparative analysis of PV infectivity more rigorous. First, our system was optimized by pseudotyping PVs with S protein mutants. Since the cytoplasmic portion of the S protein has an ER-Golgi retention signal needed for incorporation into coronaviruses that bud from endosomal membranes,^48,49^ but which may not be optimal for efficient pseudotyping of lenti- or retroviral particles that assemble predominantly at the plasma membrane, this signal should be removed. An early study on SARS-CoV by Giroglou et al.^39^ demonstrated that the C-terminally truncated spike increased PV infectivity, which led to the inclusion of this modification in the many subsequent PV systems developed for SARS-CoV-2.^6,32,41–43,50^ The truncation of last 18-19,^43,50,51^ or even 13, amino acids^41^ enhanced PV infectivity from 10 to 100 fold. Consistent with the data reported above, our study has shown that the ΔC19 mutation improved cell-free infection by 15 fold, and one-step infection by 5 fold (Figure 1D and 2B). Nevertheless, a few studies did not find a substantial influence from ΔC19^42^ or point mutations in the ER retention signal^33,52^ on PV infectivity. In agreement with published papers,^33,39,41^ we did not observe substantial differences between wt and ΔC19 S protein levels expressed on the surface of PV producing cells (Figure 1G). Thus, the mechanism of enhanced infectivity for ΔC19 PVs remains unclear, and can be related to improved incorporation of this mutant into PVs^40,41,52^ and/or stabilization of S1-S2 subunit interaction.^41^ Unlike simple truncation, the substitution of C19 with the most membrane-proximal cytoplasmic domain of gp41, ΔC19-H2^40^ did not alter PV infectivity in our tests. Crawford et al. substituted the cytoplasmic portion of the S protein with the intracellular domain from influenza hemagglutinin, and reported no improvement in PV infectivity.^33^

Unlike SARS-CoV-1 S, the S protein of SARS-CoV-2 contains a furin cleavage site (F), located a little upstream of S1/S2 boundary.^23^ It became clear early on that the presence of F increases Env-mediated cell-cell fusion, at least for *in vitro* experiments.^22,53^ However, the effects of F on virus infectivity were contradictory, i.e., either a decrease^44^ or an increase^45^ in ΔF PV infectivity relative to the wt S protein was reported. Finally, several groups found that infectivity depended on the cell target and, in particular, the entry site that the virus uses during infection.^2,22,54^ The latter is largely dependent on the S protein cleavage at S2’ site by surface protease TMPRSS2 or lysosomal cathepsins.^21^ If cells express TMPRSS2 and the virus enters via the plasma membrane, then ΔF decreased PV infection^22^; whereas TMPRSS2-negative target cells, such as the widely used 293T/ACE2 cells, were usually infected similarly^2,22^ or even better, with ΔF PVs,^44,45,52,54–56^ depending on the mutations introduced at the F site. Consistent with these reports, a 15-20 fold increase in infectivity with purified ΔF PVs in 293T/ACE2 cells was detected in this study. Strikingly, one step transfection/infection using all variants of the S protein with intact F was undetectable (Figure 2B), what was explained by massive syncytia formation induced by the S protein (Figure 2C) and by blocking inLuc-mR transduction in fused cells.^37^ Thus, measurement of cell coculture infection must adhere to the ΔF variant of spike and be used in cell-free infection for comparative purposes. Summarizing this part of the study, the ΔFΔC19 mutant of the S protein was selected as the one providing the highest sensitivity to PV infection, both in cell-free and cell coculture experimental settings.

Next, the generated PV system was validated in a neutralization test with convalescent sera. As the ΔF mutation is localized in the external part of the S protein, it can potentially influence serum neutralization activity. A comparison of the ΔFΔC19 and ΔC19 mutants revealed that ΔF required ∼ twofold higher serum concentration for PV neutralization than without ΔF (Figure 3C). This is consistent with the study by Johnson et al.,^57^ and suggests that using the ΔFΔC19 mutant spike slightly underestimated the neutralization potential of sera, but not overestimated it. Using five selected COVID-19 convalescent sera with high anti-spike titers, inhibitory activity was quantified against the ΔFΔC19 in cell-free and cell coculture modes of infection, and NT_50_ was calculated for all tested sera. It was demonstrated that all convalescent sera were at least tenfold less efficient in the neutralization of cell coculture infection, relative to inhibitory activity detected with purified PVs. Based on current literature, this is the first evidence that SARS-CoV-2 cell-to-cell infection is resistant to antibody neutralization. There is only one study in which infectivity of SARS-CoV-2 PVs was measured using a similar intron-containing reporter vector. It was based on Gaussia luciferase (Gluc) and produced minimal background activity.^43^ However, the authors of this study did not use it to measure cell coculture infection. Acknowledging that the validity of the results obtained in this study with lentiviral PVs could be heavily criticized, we conducted neutralization experiments on Vero E6 cells with live full-length SARS-CoV-2, at transmission settings that were as close as possible to those developed for the single round infection tests. The viral multiple replication assays not only confirmed the results with PVs, but also demonstrated the near complete resistance of SARS-CoV-2 cell coculture infection to neutralizing antibodies (Figure 4C). This phenomenon has been observed for a number of viruses,^58–63^ but has not been reported for coronaviruses.

Some respiratory viruses have been shown to utilize cell-to-cell transmission. Examples include induction of intercellular extensions by the influenza virus, PIV5,^64,65^ HMPV,^58,66^ and RSV^67^, usage of intercellular membrane pores by the measles virus, which is also able to infect airway epithelium.^68^ Coronaviruses extensively reorganize not only the ER-Golgi network but also change plasma membrane characteristics, inducing formation ruffles and filopodia.^69^ Ogando et al. observed that Vero E6 cells infected with SARS-CoV-2 alter their morphology by forming long filopodia with budding viruses.^70^ The study on Caco-2 cells by Bouhaddou et al. showed that the N protein of SARS-CoV-2 interacts with the kinase CK2 that leads to cytoskeleton reorganization and filopodia formation.^71^ Using scanning electron microscopy, Caldas et al. visualized thin protrusions between Vero cells with adherent virions.^72^ Whether the observed intercellular contacts play a role in SARS-CoV-2 cell-to-cell transmission *in vivo* is not known.

Massive cell-cell fusion induced by SARS-CoV-2 has been observed in cell culture^22,53^ and in human organoids,^73,74^ as well as in post-mortem material.^75–79^ However, the biological significance of syncytia formation *in vivo* remains uncertain. The most logical mechanism that would protect SARS-CoV-2 cell-to-cell transmission from antibody neutralization is a tight cellular contact with a synaptic cleft where the virus buds, and where large immunoglobulins may have difficulty penetrating. This mechanism has been implied for retroviral infectious synapses.^80^ However, such structures have not been described for SARS-CoV-2, which transmits in airway epithelium.

In conclusion, we developed lentivirus-based single round pseudoviral infection assays suitable for quantitatively measuring SARS-CoV-2 entry in cell-free and cell coculture conditions. Using this system, as well as the SARS-CoV-2 spreading assay, our study has shown that cell-to-cell infection of SARS-CoV-2 is considerably more resistant to serum neutralization than infection with purified viral particles. These results underline the importance of SARS-CoV-2 cell-to-cell transmission for virus biology, immune protection, and the development of entry inhibitors. Further experiments are required to understand the mechanisms of this resistance. The developed assays are safe, easily reproducible, and believed to be more appropriate for validating the neutralization activity of antibodies, peptides, and small molecules than the PV tests described earlier.

## Materials and methods

### Cell lines

The human embryonic kidney 293T cells were obtained through NIH AIDS Research and Reference Reagent Program. Vero E6 cells were obtained from ATCC (CRL-1586). All cell lines were cultured in high glucose Dulbecco’s modified Eagle’s medium (DMEM) (Sigma-Aldrich, USA) with sodium pyruvate, sodium bicarbonate, 10% fetal calf serum (FCS), 2 mM glutamine and 40 µg/ml gentamicin at 37°C and 5% CO2. The cells have been tested negative for mycoplasma contamination.

### Human serum samples

All serum samples were derived from the human serum biobank of the Gamaleya Center for Epidemiology and Microbiology. Study was approved by the Local ethic committee of the Moscow First Infectious Disease Hospital (Protocol #2 dated 2021-01-22).

### Plasmid construction

The plasmid pCG1-SARS-2-S coding for the codon-optimized S-protein was kindly provided by Prof. Dr. Stefan Pöhlmann (Infection biology unit of the German Primate Center, Leibniz Institute for Primate Research). C-terminal truncation of the S-protein (ΔC19), addition of 8 amino acids from the HIV gp41 (H2) and mutation of the furin cleavage site PRRA⟶A (ΔA) were introduced by PCR with Pfu polymerase (Sibenzyme, Russia) and verified by sequencing. The HIV-1 (strain NL4-3) packaging plasmids pCMV-dR8-2 (# 12263), vector pCMV-VSV-G for expression of the protein G from vesicular stomatitis virus (# 8454) were obtained from Addgene; reporter plasmids pUCHR-inLuc-mR and pUCHR-IR-GFP were described previously ^37,38^. The plasmid pUCHR-hACE2 was generated by subcloning the ACE2 coding sequence from the pCG1-hACE2 plasmid obtained from Prof. Dr. Stefan Pöhlmann (Infection biology unit of the German Primate Center, Leibniz Institute for Primate Research) into lentiviral vector pUCHR.

### Establishing 293T/ACE2 target cell line

To produce lentiviral particles, 0,35×10^6^ 293T cells were plated in 1 well of a 6-well plate in 2.5 ml of growth medium. The next day, the cells were transfected with 0.66 µg of pCMV-dR8-2, 0.88 µg of pUCHR-hACE2, and 0.22 µg pCMV-VSVG using Lipofectamine 2000 (ThermoFisher Scientific) according to the manufacturer’s instruction. At 48 h posttransfection, supernatants with pseudoviruses were cleared through 0.45 μm pore size filters and used for transduction. 8×10^4^ 293T cells per well were plated in a 24-well plate in 500 µl of growth medium. In 24 h, serially diluted lentiviral particles were added to the cells for. The percentage of ACE2-positive cells was analyzed by flow cytometry at 48 h postinfection. The sample with approximately 30% level of transduction was selected for further isolation using Sony MA900 (Sony Biotechnology, San Jose, CA, USA) cell sorter. The cells were expanded and sorted once again to enrich ACE2 positive population more than 98%.

### Generation of SARS-CoV-2 pseudotyped viral particles

2,5×10^6^ 293T cells were plated in a 10-cm dish in 10 ml of growth medium. The next day, the cells were transfected with 5 µg pCMV-dR8-2, 6.67 µg pUCHR-inLuc-mR or pUCHR-IR-GFP, and 3.33 µg of the S-protein coding plasmid using Lipofectamine 2000 (ThermoFisher Scientific) accordinhg to the manufacturer’s instruction. 48 h posttransfection, pseudoviruses were cleared through 0.45 μm filters, concentrated by centrifugation at 20000 g, 4°C, 2.5h, aliquoted and stored at −80°C. Pseudoviruses were titrated on 293T/ACE2 cells and assessed by flow cytometry (GFP) or by luciferase assay (inLuc). p24 level for each preparation was measured by the HIV p24 ELISA kit (Vector-Best, Russia).

### One step transfection/infection assay

A single-round transfection/infection test was performed in a 24 well format. 8×10^4^ 293T/ACE2 cells per well plated in 500 µl of growth medium 24 h in advance were transfected with 0.217 µg pCMV-dR8-2, 0.288 µg pUCHR-inLuc-mR, and 0.144 µg pCG1-SARS-2-S or its derivative or 0.072 µg pCMV-VSVG as a control using Lipofectamine 2000 (ThermoFisher Scientific) according to the manufacturer’s instruction. 48 h posttransfection, culture supernatants were removed, centrifuged and used for p24 calculation by ELISA. The cells were lysed with the GLO lysis buffer (# E2661, Promega), luciferase activity was determined by the Bright-Glo™ Luciferase Assay System (# E2620, Promega) using GloMax® 20/20 Luminometer (Promega).

### Detection of syncytia formation

8×10^4^ 293T cells per well were plated in a 24-well plate in 500 µl of growth medium. After 24 h, the cells were transfected with 0,5 µg pCMV-GFPt and 0,3 µg pCG1-SARS-2-S, pCG1-SARS-2-SdC19 or pCG1-SARS-2-SdC19-H2 using Lipofectamine 2000 (ThermoFisher Scientific) according to the manufacturer’s instruction. The next day, the transfected cells and 293T/ACE2 cells were detached with 1mM EDTA, mixed at the ratio of 1:1 and plated in wells of a 24-well plate with the total number of 10^5^ cells per well. Images of live cells were acquired by the Nikon eclipse Ti microscope at the x10 magnification 24 h later.

### Single-cycle cell-free infection

8×10^4^ 293T/ACE2 cells per well were plated in a 24-well plate in 400 µl of growth medium. The following day, the whole volume was replaced with 400 µl of medium containing pseudoviruses. Infection level was determined 48 h later by luciferase assay or flow cytometry. To measure neutralizing activity of sera from COVID-19 patients, sera were serially four-fold diluted in growth medium and preincubated with pseudoviruses in the total volume of 400 µl for 1 h at room temperature before addition to target cells.

### Single-cycle cell coculture infection

To generate pseudovirus-producing cells, 9×10^5^ 293T cells were plated in a 6-cm dish in 5 ml of growth medium. The next day, the cells were transfected with 1.67 µg pCMV-dR8-2, 2.22 µg pUCHR-inLuc-mR, and 1.11 µg pCG1-SARS-2-SdFdC19 using Lipofectamine 2000 (ThermoFisher Scientific) according to the manufacturer’s instruction. 24 h posttransfection, producer cells were detached with 1mM EDTA and washed twice with PBS. 2.6×10^4^ cells were mixed with serial four-fold dilutions of sera in the total volume of 200 µl and incubated for 1 h at 4°C. Next, they were mixed with 5.4×10^4^ target 293T/ACE2 cells detached with 1 mM EDTA and resuspended in 200 µl of medium. Cell mixture was plated in a 24-well plate and co-cultured in the 400 µl volume of medium. Luciferase activity was determined 48 h later.

### SARS-CoV-2 virus stock

SARS-CoV-2 strain hCoV-19/Russia/Moscow_PMVL-4 (EPI_ISL_470898) ^81^, was amplified and titrated on Vero E6 cells. Viral titers were determined as TCID_50_ by endpoint dilution assay. All experiments with live SARS-CoV-2 were performed in biosafety level 3 facility (BSL-3).

### Cell-free SARS-CoV-2 spreading assay

Vero E6 cells were plated at 8×10^4^ cells/well into 96-well plates the day prior to experiments. Serum samples were serially four-fold diluted in growth medium, mixed with MOI 0.01 of SARS-CoV-2 and incubated for 1 h at 37°C. The mixture was then added to Vero E6 cells and incubated for 5 days at 37°C. Cytopathic effect (CPE) was determined by MTT assay ^82,83^.

### Cell-to-cell SARS-CoV-2 spreading assay

Vero E6 cells were plated into T25 cell culture flask and infected with SARS-CoV-2 at MOI 0.01. The next day, infected cells were detached with Trypsin/EDTA solution (Gibco, USA), washed twice with PBS, mixed with serial four-fold dilutions of sera at 2.6×10^4^ cells per sample and incubated for 1 h at 37°C. The mixture was then combined with 5.4×10^4^ uninfected Vero E6 cells in 96-well plates in the total volume of 200 µl and incubated for 5 days at 37°C. CPE was determined by MTT assay.

### Flow cytometry

To measure S protein expression on the surface of 293T pseudovirus producing cells, 3×10^5^ live cells were incubated with the serum from a convalescent donor at the 1:100 dilution in PSB for 30 min followed by the incubation with secondary anti-human IgG antibodies conjugated with PE (1:250, # H10104, ThermoFisher Scientific) for 30 min. ACE2 expression was assessed by staining cells with polyclonal rabbit antibodies against human ACE2 (PAB886Hu01, Cloud-Clone Corp.) followed by secondary anti-rabbit antibodies conjugated with PE (1:250, # P-2771MP, ThermoFisher Scientific). Samples were analyzed on CytoFLEX S flow cytometer (Beckman Coulter). FlowJo LLC software was used for histogram visualization.

## Data analysis

The data were analyzed and visualized using GraphPad Prism 8 Software. NT_50_ values were calculated using nonlinear regression curve fit to normalized data expressed as a percentage of infectivity inhibition.

## Acknowledgements

This work was supported by the grant 075-15-2019-1661 from the Ministry of Science and Higher Education of the Russian Federation.

